# A common polymorphism that protects from cardiovascular disease increases fibronectin processing and secretion

**DOI:** 10.1101/2021.03.09.434522

**Authors:** Sébastien Soubeyrand, Paulina Lau, Majid Nikpay, Anh-Thu Dang, Ruth McPherson

## Abstract

**Background:** Fibronectin (*FN1*) is an essential regulator of homodynamic processes and tissue remodeling which has been proposed to contribute to atherosclerosis. Moreover, recent large scale genome wide association studies have linked common genetic variants within the *FN1* gene to coronary artery disease (CAD) risk.

**Methods:** Public databases were analyzed by two-Sample Mendelian Randomization. Expression constructs encoding short *FN1* reporter constructs and full-length plasma *FN1*, differing in the polymorphism, were designed and introduced in various cell models. Secreted and cellular levels were then analyzed and quantified by SDS-PAGE and fluorescence approaches. Mass spectrometry and glycosylation analyses were performed to probe possible post-transcriptional differences.

**Results:** Higher FN1 protein levels in plasma associates with a reduced risk of cardiovascular disease. Moreover, common CAD risk SNPs in the *FN1* locus associate with circulating levels of FN1. This region is shown to encompass a L15Q polymorphism within the FN1 signal peptide. The presence of the minor allele that predisposes to CAD, corresponding to the Q15 variant, alters glycosylation and reduces FN1 secretion in a direction consistent with the bioinformatic analyses.

**Conclusion:** In addition to providing novel functional evidence implicating FN1 as a protective force in cardiovascular disease, these findings demonstrate that a common variant within a secretion signal peptide regulates protein function.

## Introduction

Genome-wide Association Studies (GWAS) have identified hundreds of common single nucleotide polymorphisms (SNPs) that associate with cardiovascular disease (CAD) risk ^1–3^. Although GWAS signals are enriched for expression quantitative trait loci (eQTLs) indicating measurable impacts on transcription, the identification of causal genes is challenging since 1) the vast majority of common trait related SNPs do not overlap protein coding genes and 2) are eQTLs for multiple genes ^4,5^. Validation of statistical associations by experimental approaches is an essential first step in the development of novel therapeutic interventions. As the majority of GWAS identified variants are unlikely to be causal for several reasons, the very identification of causal SNPs among the list of GWAS identified variants is itself a complex process ^6^. Indeed, predictions place at least 80% of GWAS identified SNPs within a substantially wide 34 kbp window of causal variants in Europeans ^7^. Clearly, mechanistic insights are limited at this level of resolution, especially since *trans* (long-distance) acting variants are prevalent and may account for significant heritability ^8^. In order to pinpoint causal SNPs (“finemapping”) and identify functionally important gene targets, various approaches have been used that leverage expression data, epigenetic information, etc. ^9^. This approach has yielded surprising findings including variants located within and outside genes that regulate distal genes, as well as evidence of pervasive transcription independent mechanisms ^10–12^.

The Fibronectin 1 gene (*FN1*) encodes a group of protein isoforms that differ in sequence and localization: plasma (pFN) and cellular (cFN) ^13^. Both forms are synthesized as precursors that are processed during ER/Golgi trafficking and either enrich the local matrix environment (cFN) or secreted into the circulation (pFN) ^14^. The cellular forms exist as multiple variants that act as key structural components and regulators of the extracellular matrix (ECM), where they are deposited as insoluble fibers involved in cell adhesion. The second major form of FN1, pFN, is secreted by the liver into the circulation where it is abundant. Mice deficient in pFN display largely normal hemostasis and wound-healing, consistent with a predominant role of cFN rather than pFN in these processes ^15^. Interestingly, pFN deficient mice display increased neuronal apoptosis and larger infarct areas following focal brain ischemia, suggesting that pFN plays a protective role, possibly by activating anti-apoptotic mechanisms via integrin signaling ^15^. While pFN is not essential to vascular integrity, pFN has been shown to penetrate the vessel wall and to constitute a significant portion of arterial FN where it may participate in normal tissue remodeling, angiogenesis and neoplasia ^16–19^.

Here, we explore and clarify the mechanisms linking common GWAS identified variants that map to the *FN1*gene to CAD risk. Using bioinformatic and molecular approaches we provide evidence that differential post-transcriptional regulation underlies the *FN1*-CAD association. More specifically a polymorphism within the signal peptide of FN1 is found to regulate the ability of FN1 to be secreted. These findings provide a unique portrait of a common coding variant linked to CAD that has functional consequences at the protein level without affecting its mature amino acid sequence.

## Results

### rs1250259 links FN1 protein expression to coronary artery disease

The CAD-linked haplotype harbors several tightly linked SNPs that correlate with the disease (including top SNP rs1250229) that are causal candidates (**Table S1**). Strikingly, the region contains a single coding SNP (rs1250259), central to the CAD associated region, which was prioritized for follow-up (**Figure 1**, **Figure S1**). Interrogation of genome-wide association studies using PhenoScanner and Open Targets points to an association between the most common allele (rs1250259-A) and lower pulse pressure, reduced CAD risk, as well as to changes in blood FN1 levels (**Table S2**, **Table S3**) ^2,20–23^. While FN1 levels as a function of the rs1250259 genotype are not available, the proximal CAD protective T allele (rs1250258-T), tightly linked (R^2^ =0.99) to rs1250259-A, is associated with increased circulating FN1 and fragments thereof, suggesting that it may play a cardioprotective role (**Table S2**) ^24^.

**Figure 1.**
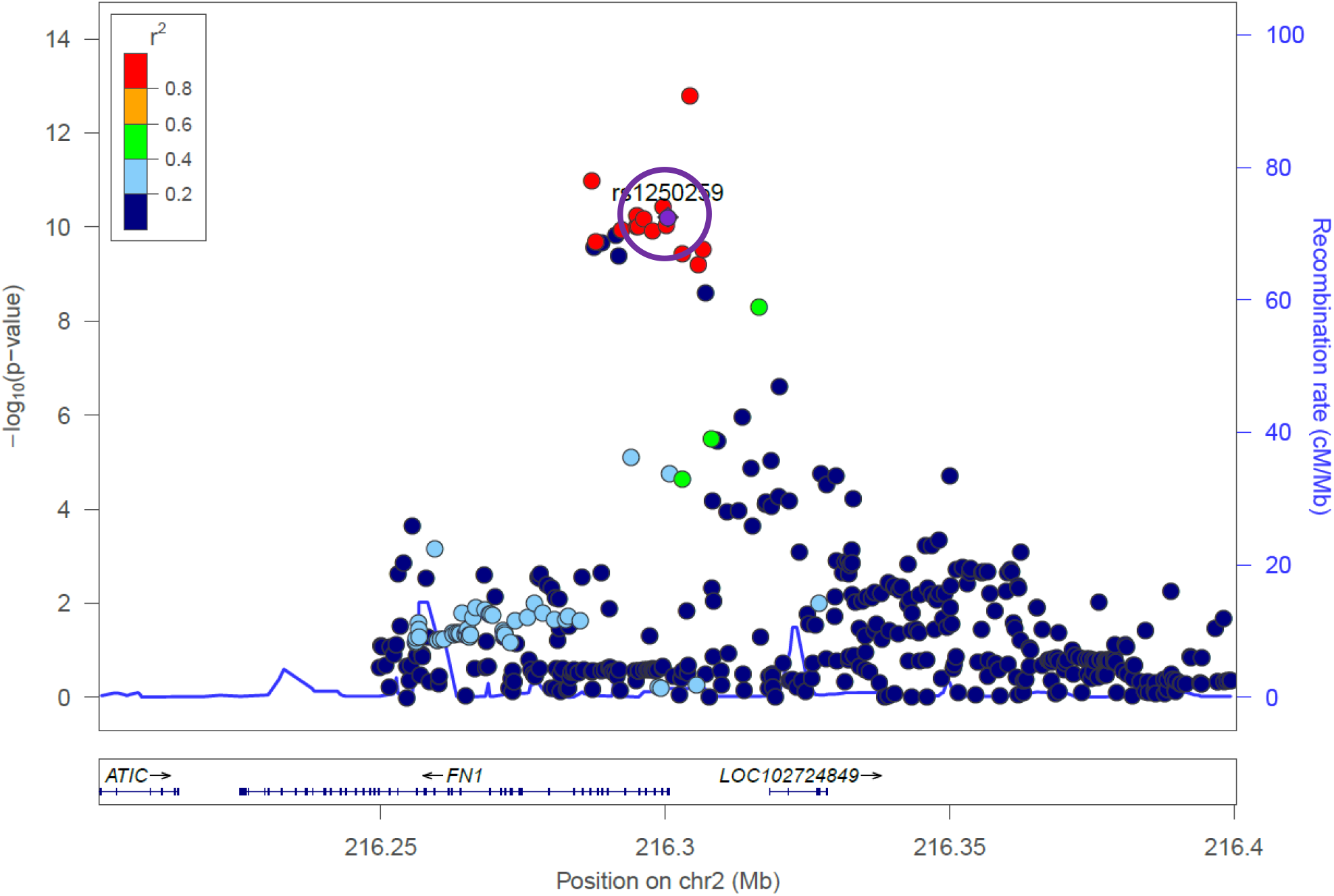
Local Manhattan plot of CAD association. CAD association data centred on rs1250259 (± 0.2 Mb), in purple and circled, from Van der Harst (https://doi.org/10.1161/CIRCRESAHA.117.312086) plotted using LocusZoom, showing a signal enrichment around the upstream region of FN1.

We next performed 2-Sample Mendelian Randomization to test a causal role for FN1 (see Materials and Methods for additional details). In this analysis, changes in FN1 protein expression linked to SNPs (whether are or not the SNPs are *a priori* correlated with CAD risk) are pooled and compared with a similar association as a function of CAD status. Consistent with individual SNP contributions, circulating FN1 (probe) and CAD were inversely correlated, with higher circulating FN1 linked to lower CAD risk (**Table 1**). Moreover, the 2-sample design supports a causal contribution of FN1, although reverse causation (CAD affecting FN1 levels) cannot be entirely excluded.

**Table 1.** Mendelian randomization reveals an impact of FN1 on CAD. Probes (protein concentration) and corresponding CAD values are from Suhre et al ^24^. Bxy, regression coefficient of x and y; se, Standard Error; p, pvalue of the beta; nsnp: number of snps used in model; Z, Z-score of the correlation. All values are rounded to 2 significant figures.

| Probe | Outcome | bxy | se | p | nsnp | Z |
| --- | --- | --- | --- | --- | --- | --- |
| 3434-34_1 | CAD | -0.059 | 0.0094 | 3.70E-10 | 5 | -6.3 |
| 4131-72_2 | CAD | -0.061 | 0.0094 | 1.00E-10 | 4 | -6.5 |

### Identification of a missense mutation within *FN1* linked to CAD that is predicted to affect secretion

Although the above analysis focused on FN1 protein expression, the contribution to *FN1* mRNA expression remained to be tested. Genotype-Tissue Expression (GTEx) data indicate that the CAD linked haplotype region (including rs1250229 and rs1250259) was not associated with statistically significant changes in *FN1* expression in any of the available tissues (data not shown). This suggests that the haplotype may affect (harder to detect) distal genes, tissues not part of the GTEx panel or a combination thereof. Alternatively, the region may affect FN1 post-transcriptionally. Translation of rs1250259 is predicted to yield protein variants harboring either a Gln (rs1250259-T) or Leu (rs1250259-A) at position 15. Of note, the SNP haplotype is defined on the positive strand while the gene is transcribed in the negative orientation (**Figure 2**). FN1 is synthesized as a precursor that undergoes removal of a ~30 amino acid region containing a hydrophobic signal peptide (which includes residue 15) and a hydrophilic short pro-sequence ^14^.

**Figure 2.**
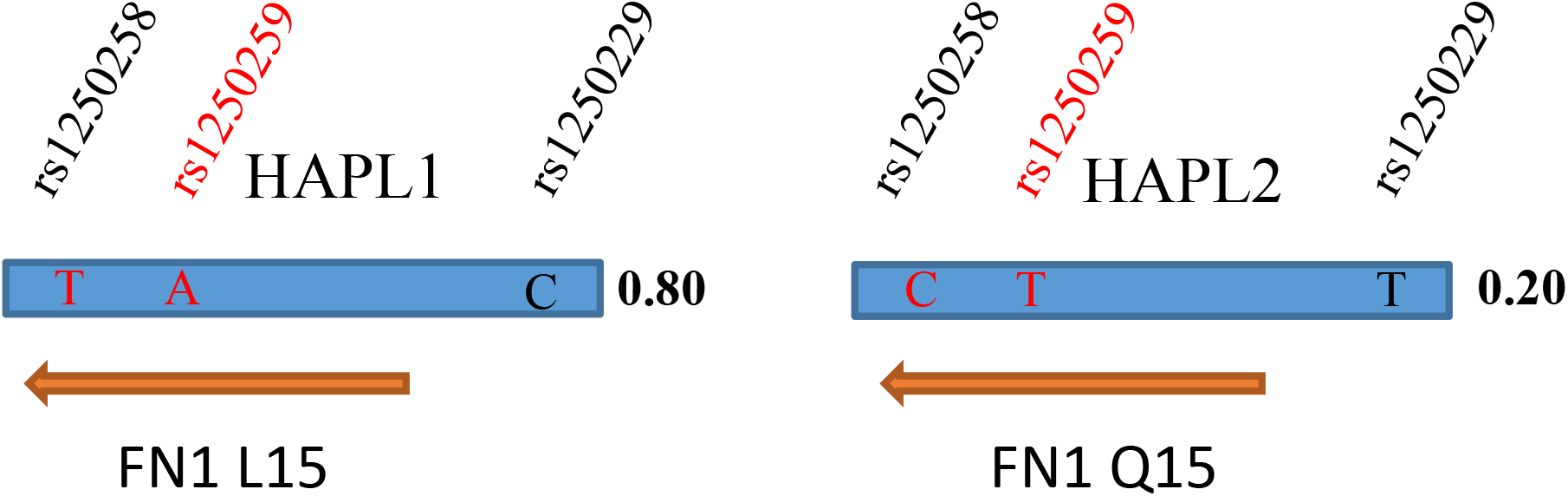
Haplotype structure around rs1250229, linked to both CAD and blood pressure. The C allele at rs1250229 correlates with the presence of rs1250259-A (r^2^=0.94), resulting in T on the coding strand of FN1 which is expressed from the negative strand. The corresponding codon (CTG) encodes a Leucine at position 15 of FN1 while the alternate codon (CAG) codes for Glutamine. Numbers on the right (Bold) are the fraction of the corresponding phased haplotype over the total number of observed haplotypes. Values are from the 1000 Genomes Project, using rs1250259 values (all populations). Genotype information for rs125058 and 59 were verified and found to be consistent with the Ottawa cohort genotyping.

### The rs1250259 affects secretion of a FN1 fusion construct in transformed and primary cell models

To examine the impact of this substitution on FN1 secretion, a model fusion protein consisting of amino acids 1-182 of FN1 fused to a GFP-HA tag moiety was generated (**Figure 3A**). This moiety is conserved in all FN1 forms and addresses technical limitations linked to the large size of FN1. The FN1 region chosen corresponds to the N-terminal heparin binding domain, which forms a well-defined region by crystallography and NMR and is shared by both secreted and cellular forms. Expression was first tested in HEK293T, a readily transfectable and widely available cell line. Following SDS-PAGE of cell lysates, a shift was observed: the Q15 form migrates slightly slower than the L15 variant (**Figure 3B**). Both fusion variants were present at comparable levels in HEK293T lysates after correcting for transfection efficiency (**Figure 3 C**).

**Figure 3.**
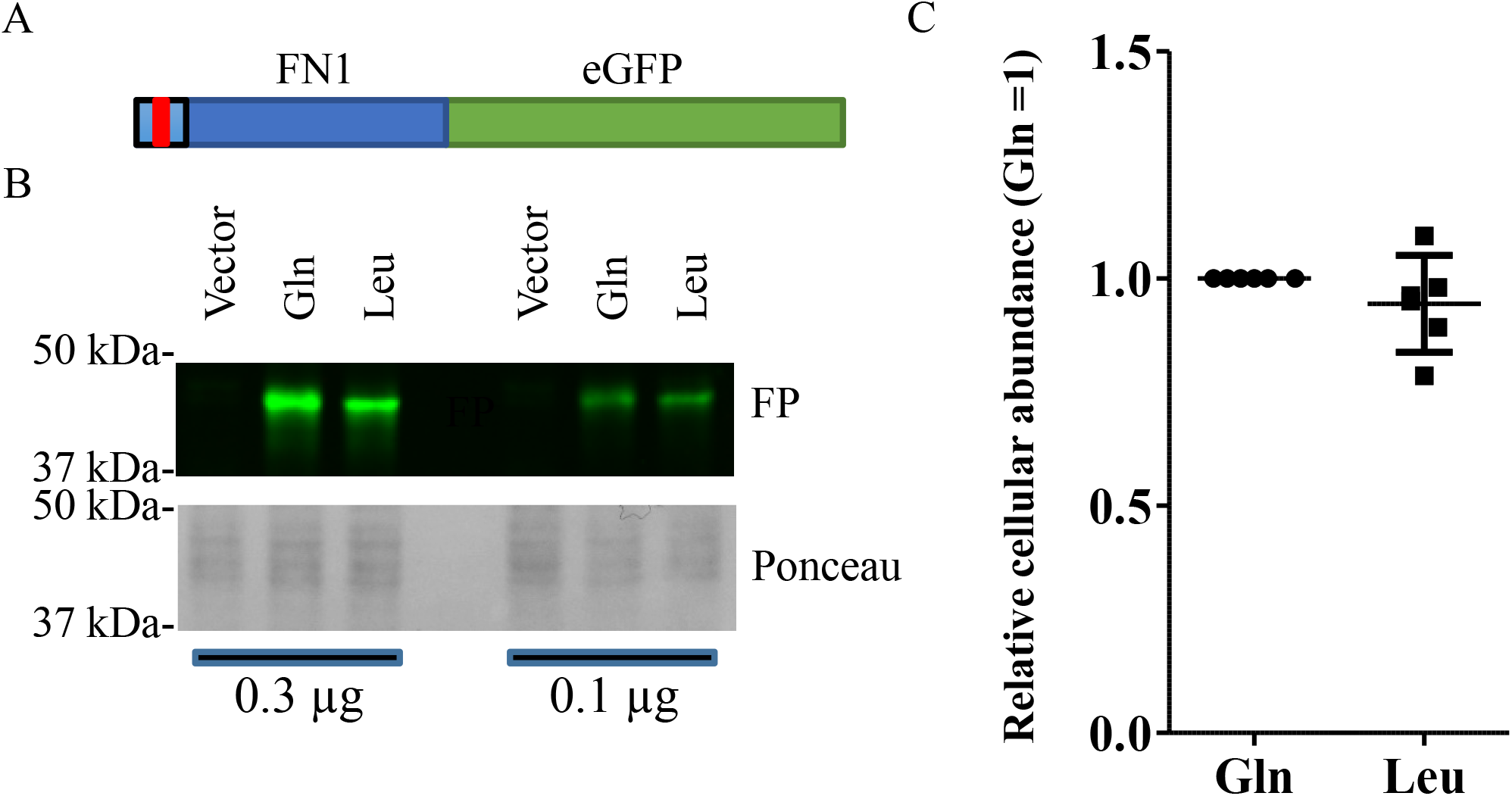
Both FN1 variants express similarly in HEK293T cells. A, Schema of the FN1-FP construct used. Drawing is approximately to scale. FN1 region corresponds to AA1-182, which encompasses, the signal peptide as well as 3 complete Fibronectin type-I domains corresponding to a previously reported crystal structure (2CG7, PDB entry). Signal peptide is in lighter blue. Red bar indicates position of the L15Q polymorphism. B,C, Analyses of cell lysates transfected with either variant (or vector alone). In B, HEK293T cells were transfected for 48 h with constructs encoding FN1-GFP fusion proteins, lysed and resolved by SDS-PAGE. The Gln variant exhibited a slight retardation relative to the Leu variant. In C, quantification of the cellular FP intensity in lysates after correction for transfection efficiency (Renilla). Each data point represents an independent experimental replicate.

Presence of the secreted protein in the media was tested next (**Figure 4**). In HEK293T and HeLa cells, transfection resulted in the secretion of FN1-GFP in the media, with the L15 exhibiting greater propensity to be secreted, defined as the signal in the culture media relative to the cellular signal. To examine the impact of this polymorphism on secretion by the liver, which is the major physiological source of pFN1, HuH-7 hepatoma cells, a widely used model of hepatocyte function, were transfected. Although the difference was smaller than observed in the epithelial models above, consistently L15 FN1-GFP was more readily secreted by HuH-7 cells. Finally, FN1-GFP was transduced into several primary cell models with relevance to CAD, i.e., adventitial fibroblasts, endothelial cells, and coronary smooth muscle cells. In all models, the Leu form was on average better secreted than the Gln form, although the difference reached statistical significance only in a lot of coronary smooth muscle cells.

**Figure 4.**
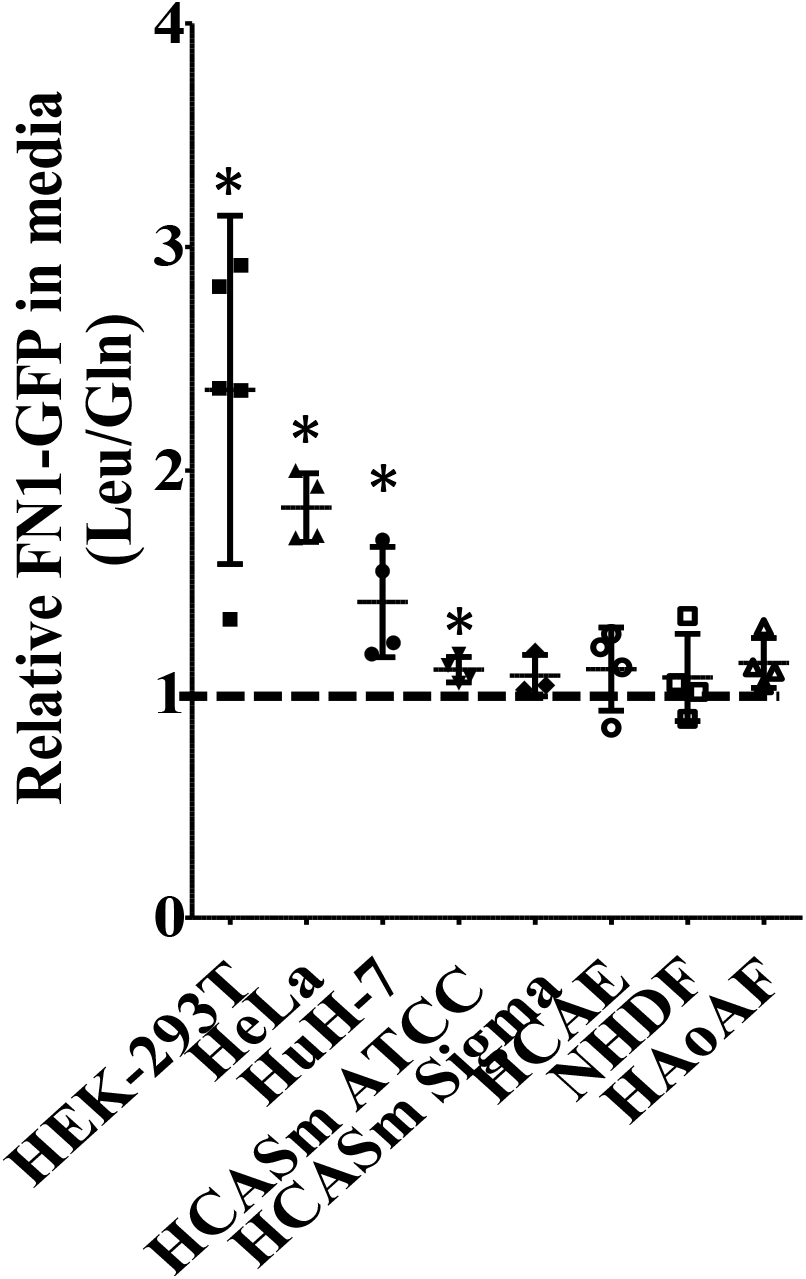
Presence of the CAD protective allele results in increased secretion of a FN1 model construct. Media and lysates from cells transduced for 48 hours with FN1-GFP plasmids encoding either Leu15 or Gln15 were analyzed by fluorometry. After correction for background signal, ratios of media to cellular GFP fluorescence were first assessed for each variant (L/Q) and the values for the Leu allele were divided by the corresponding Gln values. Values above 1 represent an enrichment of the L15 form. Results represent the mean from 3-5 independent determinations ± 95% C.I. HCASm: human coronary artery smooth muscle cells from either ATCC or Sigma; HCAE: human coronary artery endothelial cells; NHDF: Normal Human Dermal fibroblast; HAoAF: Human aorta Adventitial fibroblast.

### Secreted forms of Q15 qualitatively differ in primary cells

Examination of the variants by SDS-PAGE revealed some unexpected findings. Delivery of FN1-GFP demonstrated isoform-specific differences in the secreted forms, in a cell-type different manner (**Figure 5**). In some cells (fibroblasts, smooth muscle models as well as HeLa cells), transduction of the Q15 form led to enrichment relative to the L15 form of a slower migrating band on SDS-PAGE. By contrast, FN1-GFP from endothelial cells and HEK293T resembled HuH-7 cells in that both secreted forms exhibited qualitatively more similar profiles. Thus, in some cell types, the L15Q polymorphism appears to dictate both the quality and quantity of FN1-GFP secreted.

**Figure 5.**
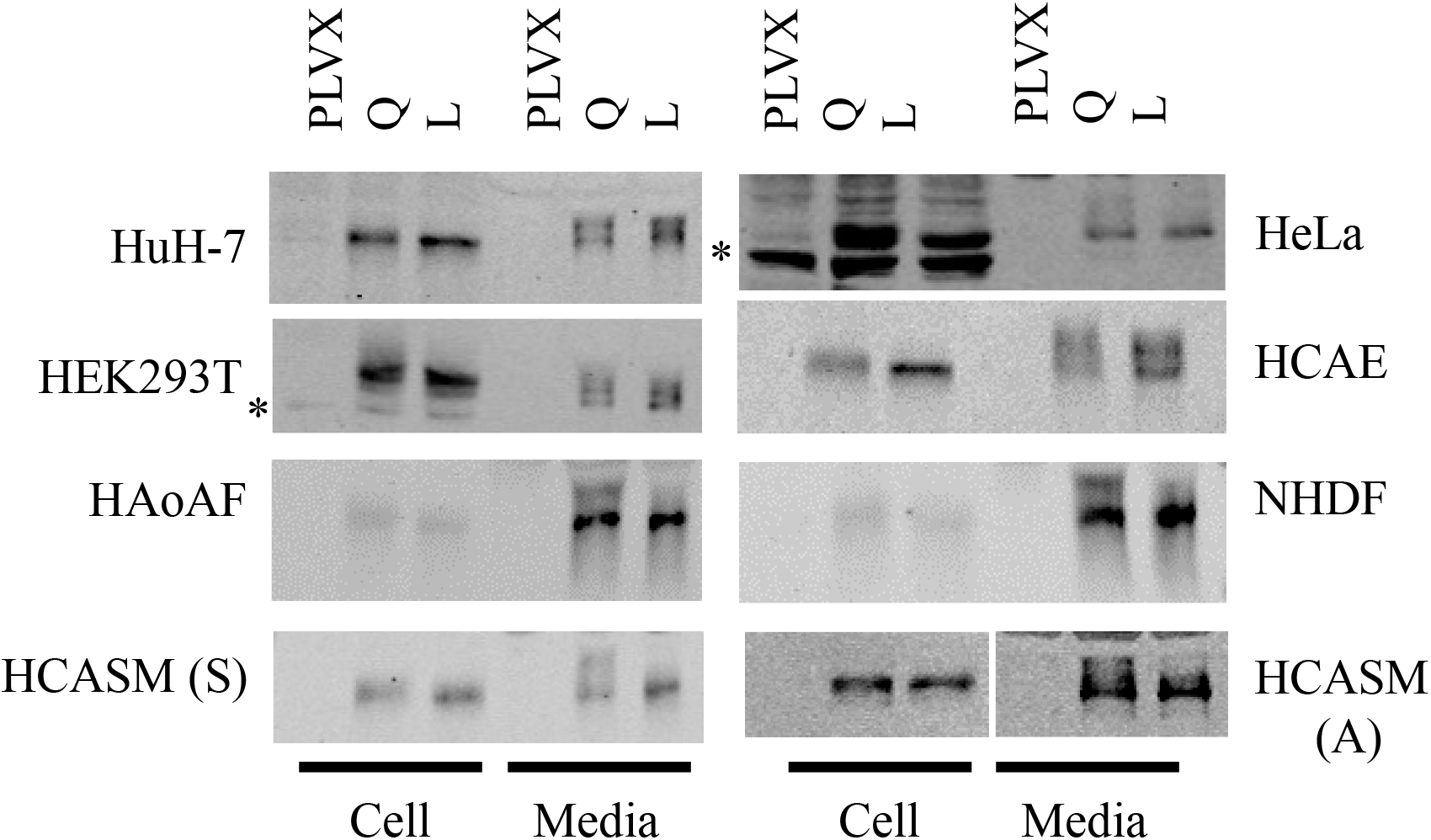
Multiple FN1-GFP species are secreted in cell cultures. Media (2% total well) and lysates (10% total well) from cells transduced for 72 hours with FN1-GFP plasmids encoding either L15 (T allele) or Q15 (A allele) were analyzed by Western blot using GFP antibodies (Sigma for all except for HeLa, Invitrogen). HCAE: human coronary artery endothelial cells; NHDF: Normal Human Dermal Fibroblast; HAoAF: Human aorta Adventitial fibroblast; HCASM: human coronary artery smooth muscle cells from either ATCC (A) or Sigma (S). Data is representative of at least 3 independent experiments. * indicates a non-specific band. Regions shown span the 37 to 50 kDa range.

### Differences in O-glycosylation account for the difference in migration

We hypothesized that this 3-5 kDa difference was due to variable levels of post-translational modification, possibly glycosylation and/or retention of pro-peptides of different lengths. As full-length pFN1 is modified post-transcriptionaly by O and N-linked glycosylation, events commonly associated with secretion, glycosylation was examined first. Both variants secreted from dermal fibroblasts were subjected to deglycosylation reactions *in vitro* using a cocktail of enzymes targeting a wide range of glycosylation chains. The incubation resulted in the disappearance of the slower migrating form (**Figure 6**). Interestingly, a longer exposure of the L15 form also shows the presence of a slower migrating band that is also sensitive to glycosylation treatment. Thus, both forms are glycosylated, albeit to different extent.

**Figure 6.**
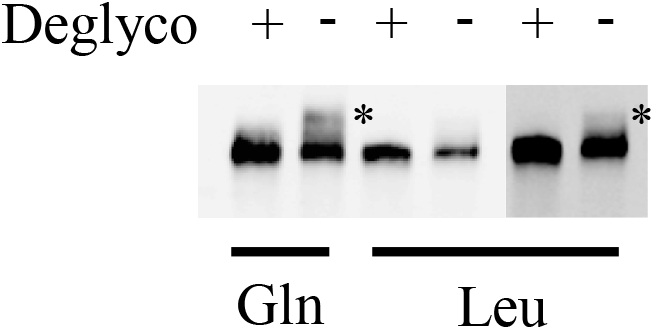
Glycosylation patterns of Q15 and L15 FN1-GFP from dermal fibroblast differ. Media from NHDF transduced for 72 hours with FN1-GFP plasmids encoding either L15 (T allele) or Q15 (A allele) were recovered by immunoprecipitation with anti-HA beads, denatured and treated with (+) or without (−) deglycosylation enzymes prior to Western blotting using an anti-GFP antibody. A higher exposure of the Leu samples is included to facilitate L-Q comparison. * indicates the position of a larger, glycosylated form. Regions shown span the 37 to 50 kDa range

The type of glycosylation involved was examined by treating cells with tunicamycin, which blocks N-glycosylation thereby interfering with protein transit through the Golgi apparatus and secretion. Inclusion of tunicamycin severely reduced the amount, and altered the migration, of full-length endogenous FN1 recovered from the media but its impact on FN1-GFP was minor (**Figure S2**). These findings point to FN1-GFP O-glycosylation exclusively.

### The L15Q polymorphism results in similar N-terminal sequences

Although the slower form reflects distinct glycosylation, the underlying cause(s) remained to be clarified. We hypothesized that distinct glycosylation profiles might result from a shift in cleavage position of the signal peptide, as suggested by SignalP (**Figure S3**). Mass spectrometry of FN1-GFP fusions isolated from the culture media of NHDF however revealed that all forms consisted of either Gln or pyroGlu at their N-termini, consistent with previous studies on full-length pFN1 ^25^ (**Figure S4**). Thus, qualitative differences in N-terminal processing are unlikely to singly account for the different glycosylation patterns. Moreover, analysis of the gel region from the L15 sample, corresponding to a putative slower form, identified the unequivocal presence of FN1-GFP of lower abundance (~ 20% of the lower form), further suggesting that glycosylation occurs on both forms albeit to different extent, with the Q15 form showing increased glycosylation.

### Quantitative differences in the secretion of the full-length FN1 variants

The impact of the L15Q polymorphism on full-length FN1 was tested next. Due to its large size, expression of a recombinant FN1 is challenging since 1) primary cells are difficult to transfect and 2) its coding sequence is too large for lentiviral delivery. For these reasons, analyses were performed on two readily transfectable cell lines: HEK293T, wherein the polymorphism had a sizeable impact on the secretion of the short FN1-GFP form and HuH-7 cells, in view of the major contribution of liver and wherein a significant but more modest impact on FN1-GFP secretion was observed. Moreover, analysis was focused on the pFN1 given its established link as a pQTL to the L15Q variant. The pFN1 construct was obtained from Addgene and modified to express a C-terminal HA tag to simplify analysis. Western blot analysis revealed an unexpected difference in HEK293T cells, in that the introduction of Q15 variant resulted in two distinct bands in cell lysates, in contrast to the L15 which showed only one (**Figure 7A**). By contrast, the forms recovered from the media were indistinguishable. Interestingly, the additional Q15 band migrated faster than the L15 band, suggesting a lower mass, that may represent an incompletely glycosylated protein. We reasoned that incomplete glycosylation might reflect a slower or impaired processing which would result in decreased secretion of mature FN1. Quantification of plasma and cellular signals indicated trends (not statistically significant) in facilitated L15 secretion (increased media signal and reduced cell signal) in both cell types **(Figure S5)**. After internal correction to cellular signal however, a clear pattern emerged whereby the L15 construct showed greater secretion, which approached statistical significance in HuH7 and reached it in HEK293T (**Figure 7B).** Thus, as for the shorter FN1-GFP, the cardioprotective L15 variant seems to facilitate secretion of sFN1.

**Figure 7.**
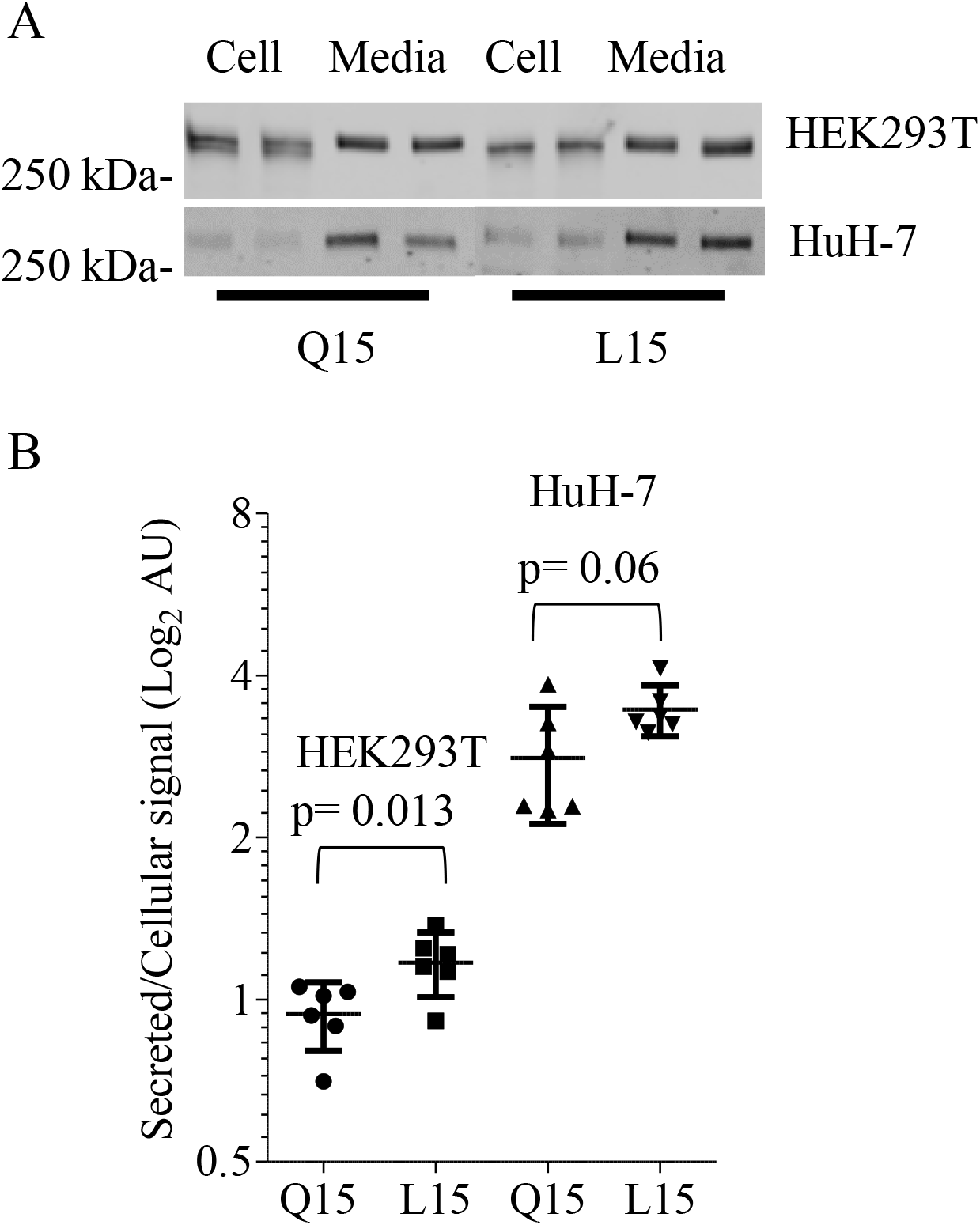
The full-length FN1 Q15 variant is less efficiently processed and secreted. Constructs encoding either variant of pFN1, tagged with a C-terminal Hemagglutinin tag, were transfected for 22 h in HEK293T or HuH-7 cells, as indicated A, Media and lysates were then analyzed by Western blot using an HA-specific antibody; no signal was detected in untransfected cells. Two independent experiments are shown for each construct. B, Quantification of FN1-HA Western blots. Signals from media (GFP) and cytosol (pRSV40 Renilla) were quantified for each transfection and data is expressed as the secreted to cellular signals for Q15 and L15 (± 95% C.I.). Each point represents a distinct experiment.

## Discussion

Here, we provide experimental evidence that the CAD protective allele of a GWAS-identified SNP increases FN1 secretion. This work provides new insights into the more global issue surrounding the role of FN1 in the pathogenesis of CAD, which is explained by genetic and environmental factors in approximately equal proportion ^26^. Moreover, it points to the role of a common variant that contributes to the heritable component of the disease. The pervasiveness of both alleles (Global Mean Allele Frequency: 0.23/0.77) in diverse ethnic groups, albeit with an uneven geographic distribution, suggests that the CAD risk variant encoding the Q15 form may confer some evolutionary benefit.

Examination of the UK biobank data via the PheWeb interface (http://pheweb.sph.umich.edu/SAIGE-UKB/gene/FN1) demonstrates an association of *FN1* with neoplasm (P=8.9e-7). While demonstrating that *FN1* may be linked to cancer, the link is with rs139452116 which results in a rare P2016L substitution and no significant association is evident for rs1250259, a SNP in strong LD with the signal peptide SNP rs1250258. This could in part account for the worsening prognosis observed previously, where strong fibronectin staining was correlated with poor survival, albeit in extreme cases only ^18^.

Signal peptides are critical for the proper maturation and secretion of extracellular proteins. Thus, mutations within secretion signal peptides can have profound repercussions if they affect the ability of the secretory apparatus to process them. A very rare R14W mutation within the signal sequence of carbonic anhydrase IV (CA4) is linked to retinitis pigmentosa and attributed to the accumulation of the immature protein within the ER, triggering the unfolded protein response and apoptosis ^27^. Unlike this extreme and rare example, the common L15Q substitution has modest quantitative and qualitative impacts on FN1. Impacts on glycosylation observed on both FN1-GFP and full-length fibronectin did not yield a coherent pattern: the Q15 form, while consistently less well secreted, exhibited increased glycosylation in FN1-GFP, at least in some primary models, but showed reduced glycosylation of full-length FN1 in our transformed cell models. Perhaps this reflects a cell-specific role of glycosylation in the control of FN1 secretion or, alternatively, a subsidiary role in defining the impact of the Q15L polymorphism. While typically associated with late protein maturation events, glycosylation can also occur co-translationally ^28,29^. Thus, we speculate that the L15Q variant may differentially affect protein translocation through the ER channel and/or peptide cleavage kinetics, resulting in altered interaction dynamics with glycosylation enzymes. For instance, slower cleavage (Q15) within the ER lumen may increase glycosylation by facilitating transient interactions with glycosylation enzymes. Additional investigations are required to resolve this question.

We demonstrate, using evidence from large datasets, that the cardioprotective allele is linked to increased FN1 secretion suggesting that circulating FN1 protects against CAD. In contrast to this hypothesis, Chiang et al employed an FN1 polymerization inhibitor to demonstrate that pFN1 blocked vascular remodelling following carotid artery ligation in a rodent model ^19^. However, the dramatic changes entailed by carotid ligation surgery in mice combined with fibronectin polymerization inhibition, which is remote from human CAD etiology. The surgical intervention notwithstanding, the effect of the inhibitor on FN1 function would greatly exceed any impact the common SNP may have on fibronectin expression. Thus, while a potent inhibition of FN1 may protect from an early stage of atherosclerosis, a modest increase may indeed be beneficial.

One limitation to our large dataset interpretation is that it is derived from an integrative analysis of distinct cohorts: a UKBiobank/- and CARDIoGRAMplusC4D meta-analysis focusing on the genetics of CAD and correlative GWAS/pQTL derived from a healthy cohort ^20,24^. However, the advantage of this approach is that by examining the impact of the SNPs predisposing to CAD on pFN1 levels in a largely healthy population, one avoids confounders frequently associated with CAD (additional underlying conditions, medications, etc.). This comes with an important limitation, as the impact of pFN1 levels on CAD is an extrapolation, albeit an informed one.

Atherosclerosis has a complex, heterogeneous etiology, involving extensive tissue remodelling characterized by smooth muscle cell proliferation which is exacerbated by hypertension as well as invasion by circulating immune cells ^30–32^. It was hypothesized that FN accumulation in the aortic media may play a role in the remodelling of the aortic wall in response to increased shear stress ^33^. This is consistent with the observation that *FN1* SNPs are also linked to blood pressure traits, suggesting that FN1 might contribute to CAD in part through the regulation of the vascular tone. Thus, the cardioprotective property of FN1 might ultimately stem from its ability to regulate vascular wall ECM assembly, by jointly affecting vascular elasticity and inflammation.

## Materials and Methods

### Tissue culture

HuH-7 were obtained from and grown in low glucose DMEM supplemented with 1 g/L glucose and penicillin (0.1 mg/ml) and streptomycin (0.1mg/ml). HEK-293T and HeLa were from the ATCC and grown in DMEM with 4.5 g/L glucose supplemented with penicillin (0.1 mg/ml) and streptomycin (0.1mg/ml). Coronary smooth muscle cells were obtained from Sigma and the ATCC and maintained in the recommended media. Human coronary adventitial fibroblasts. Normal human dermal fibroblast. human aorta adventitial fibroblasts were purchased from Lonza. All primary cells were maintained in their recommended media.

### DNA constructs

The short GFP-HA fusion proteins (L15 and Q15) were obtained by chemical synthesis of two dsDNA block variants (BioBasic) encoding amino acid 1-182 of FN1 and inserted via restriction cloning in pLVX-puro digested with EcoRI/BamHI. The construct, referred to as FN1-GFP throughout, included C-terminal GFP and HA tags. The full-length pFN1 construct was obtained from Addgene (Fibronectin-human-plasma in pMAX; Plasmid #120401 ^34^). A Q15L substitution was achieved by Q5 mutagenesis (New England Biolab) on a N-terminal Hind III/AvrII fragment transferred in pCMV5 digested similarly. Following validation by Sanger sequencing, the fragment was returned to the pMAX construct. A Hemagglutinin A epitope tag was then inserted via high fidelity assembly (NEBuilder HiFi DNA Assembly; New England Biolab) by swapping a synthetic fragment containing a C-terminal HA containing sequence within the RsrII digested pMAX pFN1 construct. The final assembly and sequences of these constructs are included in Supplemental Materials.

### Transfection and transduction

Cells were transfected with lipofectamine 3000 (ThermoFisher) using a ratio of 3:2:1 (lipofectamine 3000 (μl): P3000 reagent (μl): DNA (μg)). For infection, viral particles were first generated in HEK-293FT cells using PVLX-puro (Clontech) alongside psPAX2 and pMD2.G obtained from Addgene. Virus containing supernatants were filtered through 400 nm filters and frozen at −80 C as is. Infections were performed in the presence of polybrene (2 μg/ml).

### Secretion assays

For secretion assays, the short (FN1-GFP) construct, cells stably transduced with a pLVX FN1-GFP were grown in phenol-free media for 48-72 h prior to assay. Media was recovered while cells were rinsed in PBS and lysed in PBS/1% Triton X-100. Aliquots of cell lysates and media were then transferred to assay plates and GFP fluorescence was quantified by fluorometry on a fluorescence microplate reader (Ex 470, Em 510; BioTek). After subtraction of background values, fluorescence from media was divided by cell fluorescence. To measure FN1-GFP synthesis within HEK293T (Figure 2), FN1-GFP together with pRSV40 Renilla (2% of total transfected DNA) were transiently transfected and lysed in Passive Lysis Buffer (Promega). Cell lysates were analyzed by Western blot while Renilla luciferase activity (Glomax luminometer, Promega) was used to correct for transfection efficiency.

For the full-length FN1-HA constructs, cells (in 24 well plate format) were transiently transfected for 24 h with 0.2 ug of pMAX-FN1 and 4 ng of pRSV40 Renilla. Media were recovered while cells were rinsed in PBS and lysed in Passive Lysis Buffer (Promega). Media was directly subjected to Western Blotting, while lysates were used to normalize the expression values by measuring Renilla luciferase activity

### Immunoprecipitation and Western blotting

Cells were lysed in IP buffer (50 mM Tris-HCl, pH 7.4, 0.15 M NaCl, 0.1% Nonidet P40 (IGEPAL), 5 mM MgCl_2_) for 2 min at 4 °C. Lysates (1 mg protein equivalent) were then cleared by centrifugation (17,000 Xg) for 5 min and 20 μl of prewashed Anti-HA magnetic beads (Pierce) were added. For isolation from the media, 20 μl of beads were added to 3 ml of 400 nm filtered media harvested 72 h post-infection. Western blot was performed using 8 or 10% mini gels followed by wet transfer (1 h, 100 V) to Western grade nitrocellulose (Bio-Rad). Blots were incubated in Intercept blocking buffer (LI-COR) for 1 h and incubated for 16 h at 4 °C in the presence of cognate primary antibodies diluted 1:2000 in TBS/T (50 mM Tris-HCl, pH 7.4, 0.15 M NaCl, 0.1% Tween-20). Secondary antibodies (donkey anti-mouse (680) or -rabbit (700); LI-COR) were diluted 1:20,000. Four 1 min washes in PBS were performed after each antibody incubation.

### RNA isolation and qRT-PCR

RNA was isolated using the High Pure Isolation Kit (Roche). The Transcriptor First Strand cDNA Synthesis Kit (Roche) was used to generate cDNA using a 1:1 mixture of random hexamer and oligodT. PCR amplification and quantification were performed on a Roche LightCycler 480 using the SYBR Green I Master reaction mix (Roche). For each experiment, relative amounts of target cDNAs were first expressed relative to SRP14. Results shown represent the means of 3 biological replicates. Oligonucleotides used are described in Supplemental Materials.

### Mendelian Randomization

To investigate the possibility of an association between plasma protein level of FN1 and CAD, we did multi-SNP summary-based Mendelian randomization (MR) analysis which is also known as 2-sample Mendelian randomization ^35^. For this purpose, we obtained summary association statistics (Beta and Standard error) for SNPs (pQTLs) that are independently (r^2^<0.2) associated (P<5e^−8^) with FN1 protein level and used these as an instrument to investigate a causal effect. This means, for SNPs in our instrument (MR N_SNP_), we also obtained their summary association statistics (Beta and Standard error) with CAD and contrasted the effect sizes of the SNPs on FN1 (exposure) with the effect sizes of the SNPs on the CAD (outcome), to estimate the causal effect of FN1 on CAD. In this context, a significant negative association indicates individuals genetically susceptible to have higher levels of FN1 are at lower risk of CAD. MR analysis was done using the GSMR (*Generalised Summary-data-based Mendelian Randomisation*) algorithm implemented in GCTA software (version 1.92)^35^. As compared to other methods for 2-sample MR analysis, GSMR automatically detects and removes SNPs that have pleiotropic effect on both exposure and outcome using the HEIDI test; in addition, GSMR accounts for the sampling variance in β (beta) estimates and the linkage disequilibrium (LD) among SNPs, as such it is statistically more powerful than other 2-sample MR approaches. GSMR also incorporates a variety of quality assurance and helpful functions, notably aligning both GWAS summary datasets to the same reference allele at each SNP. Excluding SNPs that difference between their allele frequency in GWAS summary datasets and the LD reference sample is greater than 0.2, a clumping function to only keep non-correlated (r^2^<0.2) SNPs (with association P-value < 5e^−8^) in the instrument and a function to generate the scatter plot of SNP effects. Previously we used this approach to investigate the role of circulating miRNAs with regard to cardiometabolic phenotypes ^36^. We obtained GWAS summary statistics for CAD from the most recent meta-analysis of CARDIoGRAMplusC4D and UK Biobank ^20^ and GWAS summary statistics for SNPs that influence FN1 protein level from Suhre et al ^24^.

### Deglycosylation reactions

Culture media from Q15 and L15 NHDF infected for 96 h with lentiviral constructs expressing FN1-GFP were recovered, supplemented with 1 mM PMSF and centrifuged (1000 X g, 2 min) to remove cellular debris, and further cleared at high speed for 5 min (13,000 X g). Recombinant FN1-GFP was isolated from 10 ml of media (corresponding to a 10 cm culture dish) using 25 μl anti-HA Pure Proteome magnetic beads (Pierce). Beads were washed 4 × 0.5 ml of PBS/1 % Triton X-100 and resuspended in 250 μl of the same buffer. Aliquots (10%) of the isolates were used per deglycosylation reaction. Deglycosylation was performed using the Protein Deglycosylation Mix II according to the supplier’s protocol (New England Biolab); the kit includes a mixture of PNGase F, O-glycosydase and exoglycosydases to remove most glycans from target proteins. Briefly, the immunoisolated material was denatured for 10 min at 75 °C and subjected to a deglycosylation reaction for 30 min at 20 °C and 180 min at 37 C, using enzyme mix (2.5 μl) or a mock reaction (no enzyme mix) in 25 μl of bead suspension. Samples were then denatured in reducing SDS-PAGE sample buffer and analyzed by Western blotting.

### Protein Analysis by LC-MS/MS

For mass spectrometry, Q15 and L15 FN1-GFP samples were immunoprecipitated from the media of transduced NHDF as described above, resolved by reducing SDS-PAGE and stained by colloidal Coomassie blue (Simply blue); NHDF were chosen for their greater proliferative ability over coronary models while exhibiting similar shifts on SDS-PAGE. Gel pieces were than excised and destained; a gel area matching a putative, lower abundance glycosylated L form was also included, for a total of 4 samples. Two distinct biologics per sample were analyzed. Proteomics analysis was performed at the Ottawa Hospital Research Institute Proteomics Core Facility (Ottawa, Canada). Proteins were digested in-gel using trypsin (Promega) according to the method of Shevchenko ^37^. Peptide extracts were concentrated by Vacufuge (Eppendorf). LC-MS/MS was performed using a Dionex Ultimate 3000 RLSC nano HPLC (Thermo Scientific) and Orbitrap Fusion Lumos mass spectrometer (Thermo Scientific). MASCOT software version 2.6.2 ([Matrix Science](http://www.matrixscience.com/), UK) was used to infer peptide and protein identities from the mass spectra. The observed spectra were matched against custom sequences and against an in-house database of common contaminants. The results were exported to Scaffold (Proteome Software, USA) for further validation and viewing.

### Statistical analysis

To estimate statistical significance of experimental findings, unpaired Student t-test (2-tailed) were performed in GraphPad Prism.

## Supporting information

Supplemental Tablles

## Acknowledgements and Funding

This work was funded by a Canadian Institutes for Health Research Foundation grant (FRN:154308; RM).

