## Supplemental figures for "A common polymorphism that protects from cardiovascular disease increases fibronectin processing and secretion"

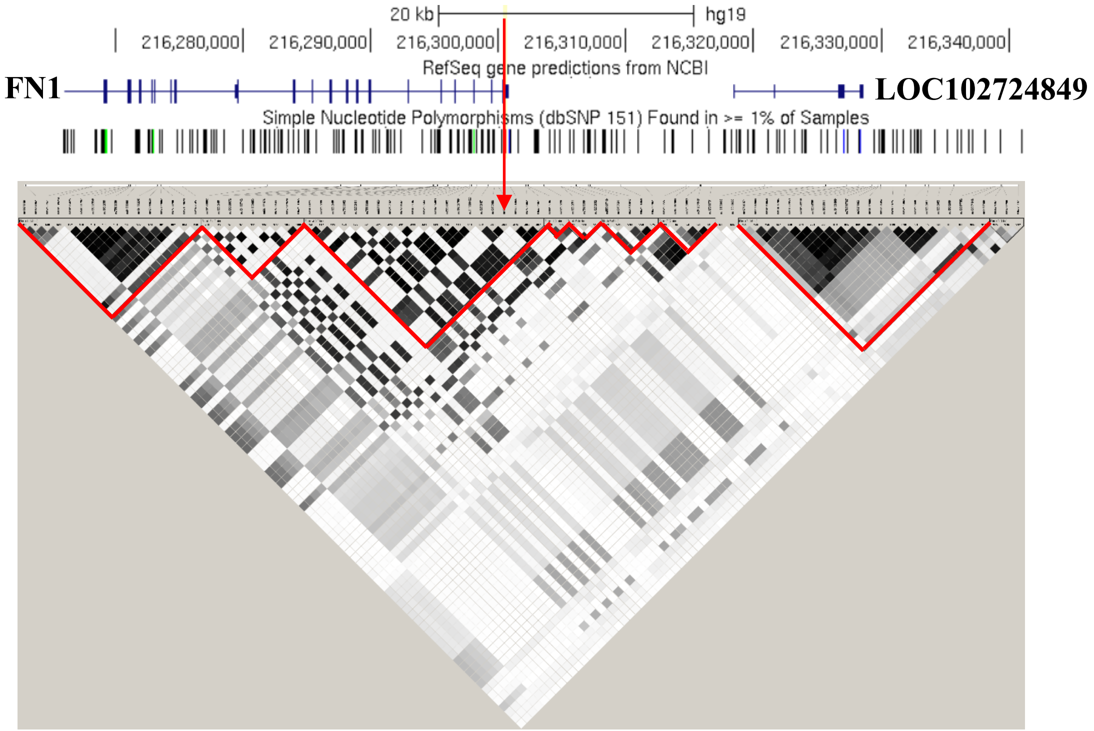


**Fig S1. Haploview map with overlapping genes**. Region spanning rs1975319 to rs6726337 is shown. LD intensity is proportional to linkage ($R^2$) values. Blocks were defined using the LD spine method. LD Only common SNPs (frequency > 0.05) are shown. Red arrow points to rs1250259.

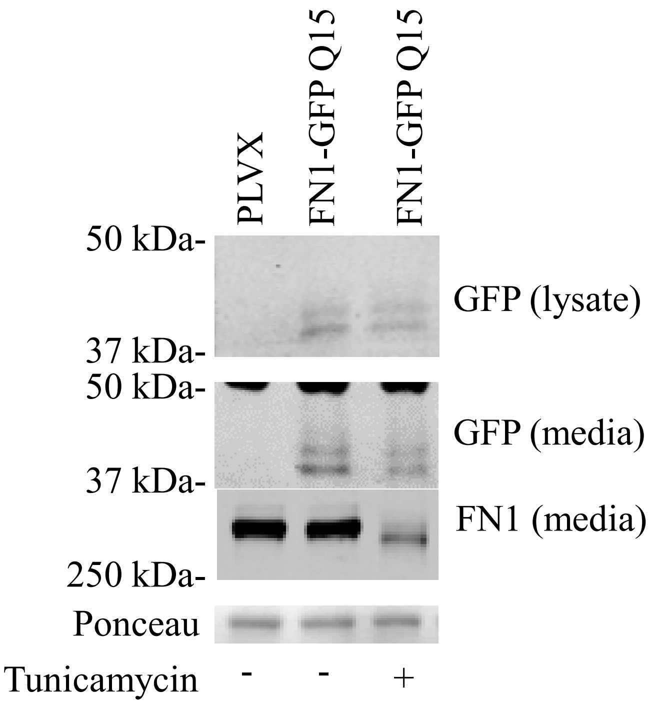


**Fig S2**. **Secreted FN1-GFP is insensitive to Tunicamycin.** HuH-7 cells stably transduced with FN1-GFP were treated for 24 h with 10 µg/ml tunicamycin. Media were harvested and analyzed by Western blotting on a 8% SDS-PAGE gel for FN1-GFP in lysates and media and endogenous media full-length FN1, as a control for Tunicamycin efficacy. Ponceau stain of a ~ 150 kDa section matching the media samples is included as a loading control.

A

| | Signal peptide prediction | Cleavage site | Probability of cleavage |
|---|---|---|---|
| L | 0.99 | TGA-SK (POS 26-27) | 0.27 |
| Q | 0.97 | GTA-VP (POS 20-21) | 0.26 |

B

| | Cleavage site | Probability of cleavage |
|---|---|---|
| L | STG-AS (POS 25-26) | 0.32 |
| Q | STG-AS (POS 25-26) | 0.27 |

**Fig S3. Bioinformatic predictions of L15Q variations on signal peptide cleavage**. The N-terminal domain of FN1 was analyzed via SignalP -5.0 (http://www.cbs.dtu.dk/services/SignalP/) (**A**) or TargetP-2.0 (http://www.cbs.dtu.dk/services/TargetP/) (**B**) to predict the impact of the Q15L natural variant on processing. Both approaches predict modestly reduced cleavage for the Q15 variant but only SignalP predicts a shift in the cleavage position.

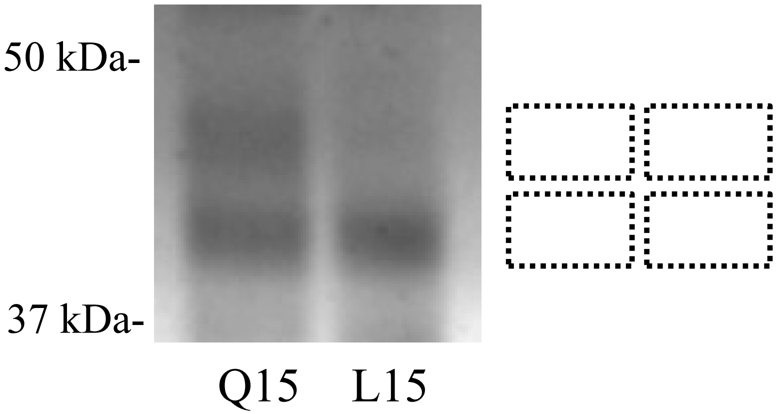


**Fig S4**. **Representative Coomassie stain of an SDS-PAGE gel of immunoprecipitated FN1-GFP.** Q15 and L15 fusion proteins were isolated from 2.5 ml of media and analyzed by Western blot. Gel pieces derived from the slower (glycosylated) and faster forms were analyzed separately by LC-MS/MS. The boxes indicate the corresponding regions isolated from the gel (for clarity shown on the side of the gel).

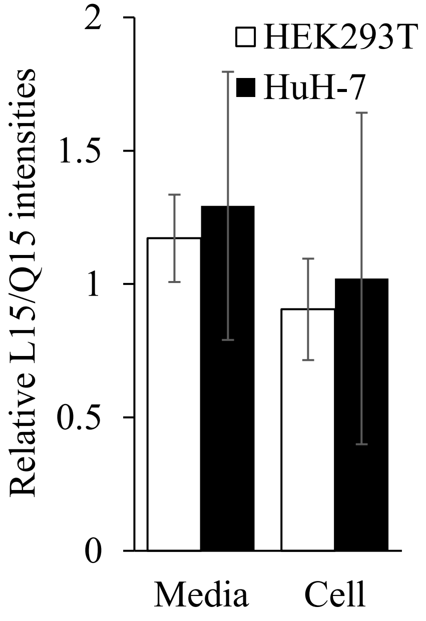


**Fig S5. Quantification of L15 and Q15 variants in the media and cell.** Data is expressed as the ratio of the L to Q Western blot signals in each compartment. Differences were not statistically significant.
